# Reduced inhibition in depression impairs stimulus processing in human cortical microcircuits

**DOI:** 10.1101/2021.02.17.431698

**Authors:** Heng Kang Yao, Alexandre Guet-McCreight, Frank Mazza, Homeira Moradi Chameh, Thomas D. Prevot, John Griffiths, Shreejoy J. Tripathy, Taufik A. Valiante, Etienne Sibille, Etay Hay

**Author notes:** These authors contributed equally (Heng Kang Yao, Alexandre Guet-McCreight). Corresponding author: Dr. Etay Hay, Krembil Centre for Neuroinformatics, Centre for Addiction and Mental Health, 250 College St, Toronto, Ontario, M5T 1R8.

## Abstract

Cortical processing depends on finely-tuned excitatory and inhibitory connections in neuronal microcircuits. Reduced inhibition by somatostatin-expressing interneurons is a key component of altered inhibition associated with treatment-resistant major depressive disorder (depression), which is implicated in cognitive deficits and rumination, but the link remains to be better established mechanistically in humans. Here, we tested the impact of reduced somatostatin interneuron mediated inhibition on cortical processing in human neuronal microcircuits using a data-driven computational approach. We integrated human cellular, circuit and gene-expression data to generate detailed models of human cortical microcircuits in health and depression. We simulated microcircuit baseline and response activity and found reduced signal-to-noise ratio and increased false/failed detection of stimuli due to a higher baseline activity in depression. Our results thus applied novel models of human cortical microcircuits to demonstrate mechanistically how reduced inhibition impairs cortical processing in depression, providing quantitative links between altered inhibition and cognitive deficits.

## Introduction

Cortical processing relies on specific interactions between different types of neurons with distinct electrical properties and synaptic connectivity^1,2^. Accordingly, there is increasing evidence that cortical dysfunction involves changes in the cellular and microcircuit properties. Altered cortical inhibition is implicated in a variety of brain disorders such as autism, schizophrenia and major depressive disorder (depression)^3–8^. Reduced dendritic inhibition from somatostatin (SST) interneurons is a key component of the altered inhibition that is associated with treatment-resistant depression and several other disorders^3,4,7–11^. Recent findings showed a significantly lower SST expression by SST interneurons in post-mortem cortical tissue from depression patients, which indicates a reduction in SST interneuron mediated inhibition in depression^11^. Accordingly, silencing SST interneuron inhibition in rodents produced anxiety and depression symptoms, and new pharmacology facilitating SST interneuron inhibition through alpha-5-gamma-aminobutyric-acid-A (α5-GABA_A_) receptors led to pro-cognitive and antidepressant effects^10,12^. However, the link between reduced SST interneuron inhibition and cortical deficits remains to be better established mechanistically, particularly in humans.

In the cortex, SST interneurons primarily target the apical dendrites of pyramidal (Pyr) neurons and provide synaptic and extrasynaptic (tonic) inhibition through activation of α5-GABA_A_ receptors^2,13^. SST interneurons mediate lateral inhibition of Pyr neurons through inhibitory disynaptic loops, where facilitating excitation from a Pyr neuron is sufficient for triggering spikes in an SST interneuron and consequently inhibit neighbouring Pyr neurons^14,15^. At the microcircuit level, SST interneurons are involved in maintaining a low baseline activity of Pyr neurons, but are largely silent during the early response of the cortical microcircuit to stimuli^1^. The role of SST interneurons in baseline activity was further supported by a two-fold increase in the baseline firing rate of Pyr neurons when SST interneurons were silenced^1^.

In line with these findings, a leading hypothesis suggests that a reduced SST interneuron inhibition in depression would increase the baseline cortical activity (noise) while minimally impacting stimulus response (signal) and thus decrease the signal to noise ratio (SNR) of cortical processing^4^. This is further supported by studies in animal models of depression, where positive allosteric modulation of α5-GABA_A_ receptors (which are targeted by SST interneurons) led to pro-cognitive effects and recovery from depression symptoms in rodents^10,12^. However, it remains unclear whether the level of reduced SST interneuron inhibition estimated from gene expression data in human depression would have a significant effect on baseline activity and thus on cortical processing. In addition, the effect of reduced SST interneuron inhibition on Pyr neuron baseline and response activity is difficult to predict due to the inter-connectivity between the different neuron types in the microcircuit. Moreover, the implications of reduced inhibition on processing SNR and signal detection in depression remain to be determined, to better link the cellular and circuit effects to specific cognitive deficits in depression such as patients being less able to identify relevant signals from noise^16^, or unable to suppress loops of internal thoughts in rumination^3^.

The link between reduced SST interneuron inhibition and depression is supported by pre-clinical animal models^7,10,12^, but there is a need for integrative mechanistic studies to assess whether this translates to humans. There are cellular and circuit similarities between rodents and humans^17^, e.g. in the intrinsic firing properties and connectivity patterns of cell types, but also important differences. Human inhibitory synapses from SST and parvalbumin (PV) interneurons onto Pyr neurons are stronger, with lower probability of synaptic failures and larger postsynaptic potential (PSP) amplitudes compared to rodents^14,18,19^. The connection probability between Pyr neurons in human layer 2/3 (L2/3) is also significantly higher than in rodents^20^. Furthermore, human Pyr neuron and interneuron morphologies are larger than in rodents, with longer and more complex dendrites allowing for more complex signal integration^21,22^, and a more compartmentalized apical dendritic tree^23^. As such, it is imperative to study the mechanisms of dysfunctional cortical processing in depression in the context of human microcircuits.

Computational models are well suited to bridge this gap, due to limitations in monitoring human neuron types and microcircuits *in vivo* and due to the increasing data availability of neuronal and synaptic recordings in healthy human cortical slices resected during surgery, especially from cortical L2/3^14,20,24^. These novel data can be integrated into detailed models of human cortical microcircuits to study human cortical processing mechanistically in health and disease. Available models of human L2/3 neurons reproduced some of the firing properties^25^, but only partly reproduced the frequency-input relationship. These models were not constrained with sag-current properties, which depend on dendritic h-current, and thus were limited in capturing dendritic input integration properties of the neurons. In addition, there are currently no models for human synaptic connections and cortical microcircuits. Therefore, there remains a need to integrate the available human electrophysiological data into detailed cortical microcircuit models, and their baseline and response activity.

In this study, we tested whether the reduced SST interneuron inhibition estimated from gene expression in depression results in significant changes in the SNR of human cortical processing and in the quality of stimulus detection. Using a computational approach, we integrated human cellular, synaptic and gene expression data from recent studies to generate detailed models of human cortical L2/3 microcircuits that included the major neuron types with their firing and input integration properties, synaptic models, and connection probabilities. We generated depression microcircuit models by reducing the SST interneuron synaptic and tonic inhibition in the circuit according to gene expression data from recent human studies^11^. We simulated baseline activity and stimulus response as constrained by previous studies of neuron-type activity profiles, to characterize the SNR of cortical processing and the failed/false detection rates of stimuli in health and depression.

## Results

We first generated data driven models of human cortical microcircuits and their baseline and response activity, using human cellular and circuit data whenever available, and data from rodents otherwise (**Table S1**). The process included modeling single neurons to represent each of the four key neuron types, modeling synaptic properties, and modeling microcircuits with the appropriate proportion of the different neuron types and the connectivity statistics.

### Human cortical L2/3 neuron and synaptic models reproduce experimental properties

We generated single neuron models of the four major cell types in cortical L2/3, using genetic algorithm optimization, to reproduce their electrical properties as measured in human cortical slices. The features of the Pyr neuron model firing in response to depolarizing step currents (e.g. spike rate, height, half-width, adaptation), the frequency-input curve, and sag voltage in response to hyperpolarizing current steps were all within the range (1 - 2 SD) of the experimental population (**Fig. 1a, Table S2)**, except for the after-hyperpolarization depth which was marginal (4 SD). The passive and active firing features of the SST, PV and vasoactive intestinal peptide (VIP) interneuron models matched the values of the corresponding human neurons (**Fig. 1b - d, Tables S3 – S5**), and were also within the experimental variance of population data from corresponding neurons in rodents^26–28^. Spike half-width in the PV models was further from the experimental variance (4.4 - 4.8 SD), which is commonly the case due to the fixed kinetics of the underlying channel models^25,29^.

**Figure 1.**
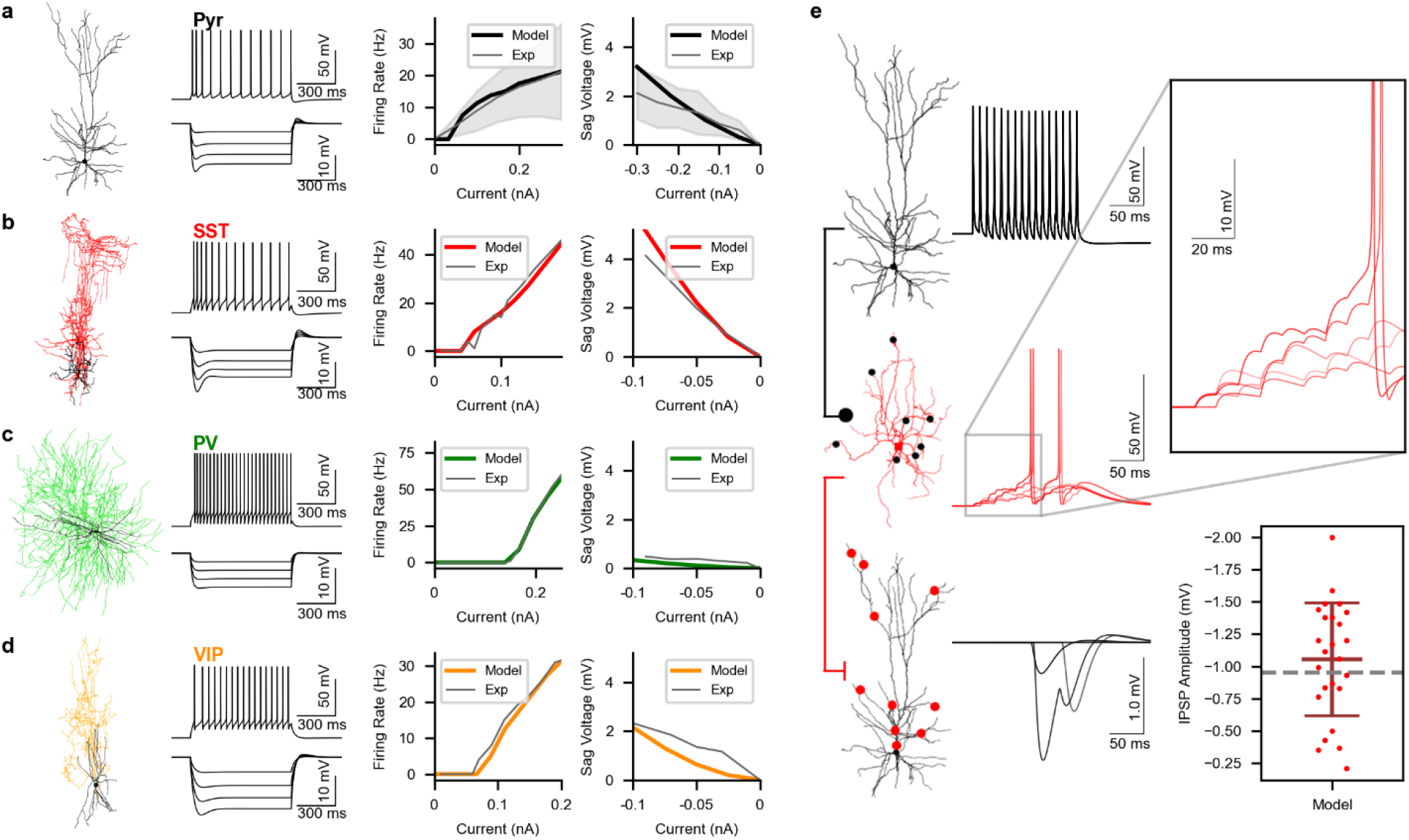
Model human cortical neurons and synaptic connections reproduce experimental properties. **a.** Left - reconstructed human L2/3 Pyr neuron morphology. Middle - Pyr neuron model response to depolarizing and hyperpolarizing step currents. Right - firing frequency-input curve and sag voltage-input curve of the Pyr neuron model (black) were within the experimental population range (gray, n = 28 neurons). **b - d.** Same as a, but for SST (b), PV (c) and VIP (d) interneurons. Firing frequency-input curve and sag voltage-input curve of the interneuron models (SST, PV, VIP: red, green, orange) matched experimental values of the corresponding single neurons (gray). **e.** Simulated voltage traces of a human disynaptic inhibition loop model. A Pyr neuron (top) fired 15 spikes at 100 Hz, and the resulting EPSP summation (inset) in an SST interneuron (middle) triggered two spikes, which elicited IPSPs in another Pyr neuron (bottom). The dots on the morphologies indicate synaptic contact locations (black: excitatory; red: inhibitory). The graph on the bottom right shows the simulated IPSP amplitude between SST and Pyr neurons (n = 25 random connections) compared to experimental value (dashed line^14^).

Next, we generated models of synaptic connections between the neuron types. We constrained the connections between Pyr neurons and SST interneurons to reproduce properties of the disynaptic inhibition loop as recorded in human brain slices (**Fig. 1e**). The model synaptic connections from Pyr neurons onto SST interneurons elicited facilitating excitatory PSPs (EPSPs) with amplitudes as seen experimentally^14^ (model: 2.16 ± 1.35 mV, experimental: 2 mV) and were sufficiently strong to trigger 1 - 2 spikes in the SST interneuron in response to a train of inputs from a single Pyr neuron. The resulting SST interneuron inhibitory PSP (IPSP) amplitudes in a neighbouring Pyr neuron agreed with the experimental value^14^ (model: −1.05 ± 0.44 mV, experimental: −0.95 mV). In addition, we constrained the EPSP amplitudes between Pyr neurons, and the connections between Pyr and PV neurons to reproduce the experimental values seen in human neurons (Pyr→Pyr model: 0.42 ±0.37 mV, experimental: 0.42 ±0.45 mV; Pyr→PV model: 3.29 ± 1.16 mV, experimental: 3.29 ± 1.12 mV; PV→Pyr model: −2.23 ± 1.2 mV, experimental: −2.23 ± 1.0 mV). The remaining types of synapses, primarily involving VIP neurons, were constrained using rodent data since human data was unavailable (**Table S7**). In addition to synaptic inhibition, we modeled tonic inhibition using voltage-clamp recordings of the tonic current in human pyramidal neurons (see Methods). We estimated the tonic conductance using the measured difference between tonic inhibition current and when GABA_A_ receptors were blocked. We reproduced the target tonic inhibition current amplitude with a tonic conductance of 0.938 mS/cm^2^ and applied it to all neurons^30^.

### Increased baseline activity (noise) in depression microcircuit models

We used the neuron and synaptic connection models to simulate human cortical L2/3 microcircuits of 1000 neurons, with the experimental proportions of the different neuron types, their connectivity and column dimensions (**Fig. 2a - b**). The neurons received random background excitatory input corresponding to cortical and thalamic drive, to enable recurrent activity. We tuned the background excitatory input level and the microcircuit connection probabilities between neuron types to reproduce the baseline firing rates previously reported for the neuron types *in vivo* (**Fig. 2c - d, f**). The average firing rate for simulated Pyr neurons was 1.10 ± 0.05 Hz, PV: 10.31 ± 0.50 Hz, SST: 5.64 ± 0.18 Hz and VIP: 3.51 ± 0.32 Hz (n = 10 microcircuits, rates for non-silent neurons as in experiments, see Methods). In addition, we have constrained the microcircuit to reproduce the effects of SST interneuron silencing, which has previously been reported to result in a doubling of baseline Pyr neuron spike rates^1^ (**Fig. 2e**, 1.10 ± 0.05 Hz vs. 2.08 ± 0.08 Hz, t-test, *p* < 0.001).

**Figure 2.**
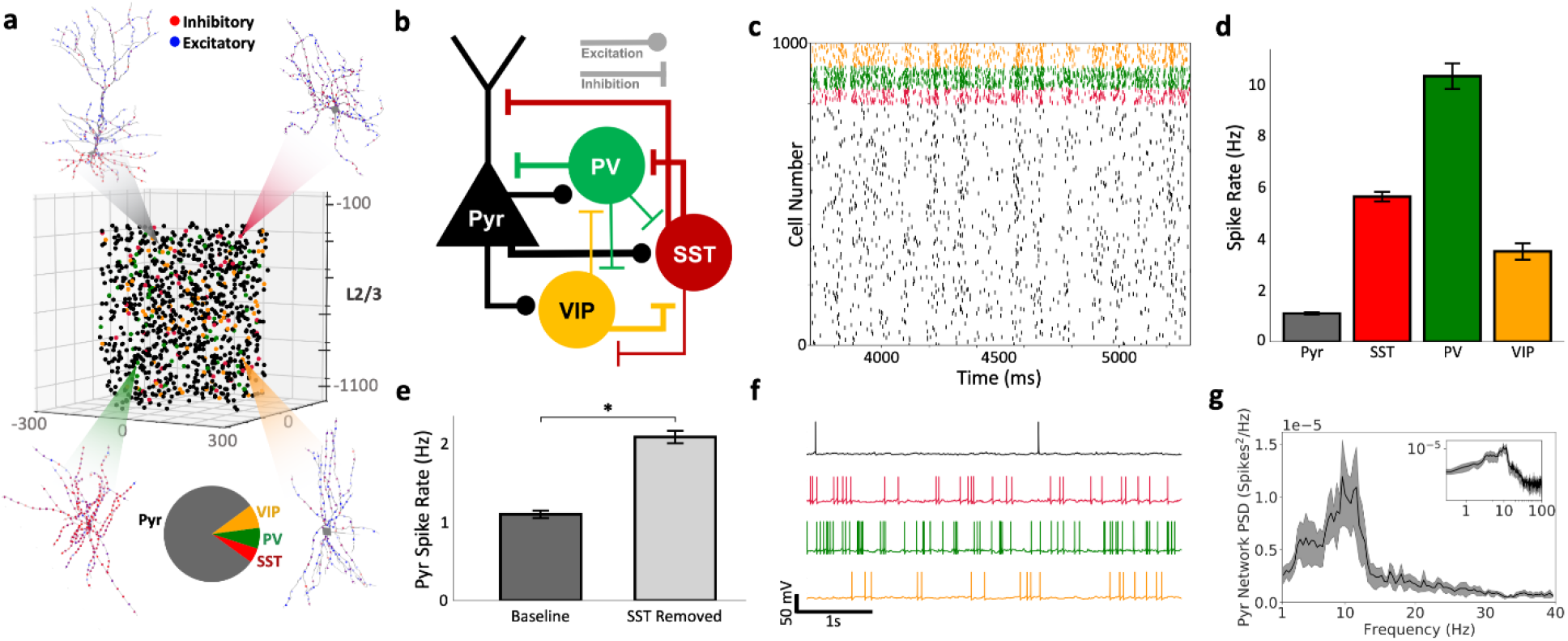
Detailed models of human cortical L2/3 microcircuits reproduce features of intrinsic circuit activity. **a)** The model microcircuit comprised of 1000 neurons, with the somas distributed in a 500×500×950 μm^3^ volume along layer 2/3 (250 to 1200 μm below pia). The proportions of the different neuron types were based on the experimental literature (pie chart, Pyr: 80%; SST: 5%; PV: 7%; VIP: 8%). The neurons were modeled with detailed morphologies, as shown in Fig 1, and connected according to the experimental statistics (see Methods). The blue and red dots on each morphology denote example excitatory and inhibitory synapses, respectively. **b)** The schematic connectivity diagram highlights the key connections between different neuron types in the microcircuit. **c)** Example raster plot of spiking in different neurons in the microcircuit, color-coded according to neuron type. Neurons received background excitatory inputs to generate intrinsic circuit activity. **d)** Spike rates in the different neuron types reproduced experimental baseline firing rates (mean and standard deviation, n = 10 randomized simulated microcircuits). **e)** A significant 2-fold increase in mean Pyr neuron spike rate when SST interneurons were silenced reproduced experimental results (n = 10 microcircuits, paired-sample t-test; *p* < 0.001). **f)** Example simulated voltage traces for each neuron type. **g)** Spikes PSD of Pyr neurons, bootstrapped mean and 95% confidence intervals (n =10 randomized microcircuits). Inset – PSD in log scale, illustrating the 1/f relationship between power and log frequency.

The microcircuit models had an emergent property of oscillatory activity, which was not constrained for explicitly, where population spiking oscillated primarily in the theta (4 – 8 Hz) and alpha (8 – 12 Hz) frequency bands, and on average exhibited peak frequencies around 10 Hz (**Fig. 2g**). The power spectrum density (PSD) plot also exhibited a 1/f relationship between power and log frequency (**Fig. 2g**, inset). These oscillatory properties closely agree with the oscillations seen in human cortical signals *in-vitro* and *in-vivo*^31–34^, providing a validation in support of the models capturing key properties of human cortical L2/3 microcircuits.

We next modelled human depression microcircuits by reducing SST interneuron synaptic and tonic inhibition conductance onto the different neuron types by 40%, according to post-mortem gene expression data in depression (**Fig. 3a - b**). We then compared the baseline activity in healthy and depression microcircuits, across all neurons of each type. The average baseline firing rate of Pyr neurons was significantly higher in depression microcircuits compared to healthy microcircuits (**Fig. 3c - d**, healthy: 0.77 ± 0.05 Hz, depression: 1.20 ± 0.07 Hz, n = 200 randomized microcircuits, t-test *p* < 0.05, Cohen’s *d* = 7.06). Next, we quantified the effect of different levels of SST interneuron inhibition reduction on baseline rates of Pyr neurons by simulating microcircuits with 0 - 100% reduction compared to the healthy level. The baseline firing of Pyr neurons increased approximately linearly with reduced SST interneuron inhibition (**Fig. 3d**). Similar increases in firing rates were observed in the interneuron populations of the depression microcircuits compared to healthy microcircuits (**Fig. 3e**, SST: 5.62 ± 0.27 vs. 7.53 ±0.39 Hz; PV: 10.19 ± 0.51 vs. 15.99 ± 0.54 Hz; VIP: 3.52 ± 0.37 vs. 7.26 ± 0.49 Hz, n = 200, *p* < 0.05 for all). Interneuron rate increase was thus largest in VIP interneurons (106%) and PV interneurons (57%), and more moderate in SST interneurons (34%). The higher activity in depression microcircuits when no stimulus was given therefore indicated an increased noise level with respect to processing incoming stimuli. To dissect the individual contribution of reducing synaptic vs. tonic inhibition, we repeated the simulation with selectively reducing SST interneuron synaptic inhibition or reducing SST interneuron tonic inhibition. We found similar levels of increase in baseline firing rates in each condition (synaptic only: 0.98 ± 0.06 Hz; tonic only: 0.98 ± 0.06 Hz). The effect of the joint synaptic and tonic inhibition reduction was therefore approximately the sum of the separate effects (separate: 1.19 ± 0.08 Hz, joint: 1.20 ± 0.07 Hz, Cohen’s *d* = 0.13).

**Figure 3.**
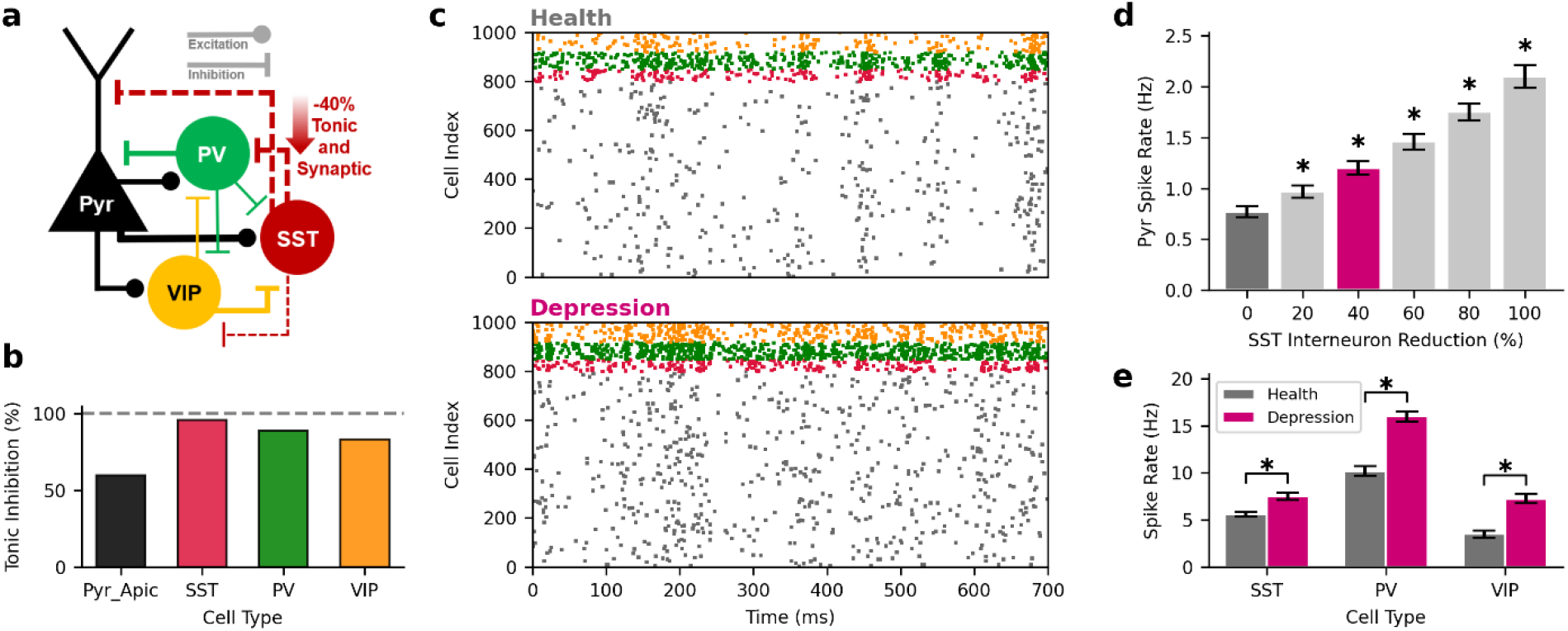
Increased baseline activity (noise) in depression microcircuit models. **a.** Depression microcircuits were modeled according to gene expression data, with −40% synaptic and tonic inhibition conductance from SST interneurons onto the other neuron types. **b.** Relative tonic inhibition conductance onto the different neuron types in depression microcircuits, compared to healthy microcircuits (dashed line). **c.** Example raster plot of simulated baseline spiking in a healthy microcircuit model (top) and depression microcircuit model (bottom). **d.** Increased intrinsic Pyr neuron firing in depression microcircuit models (n = 200 randomized microcircuits per condition, purple - depression, *p* < 0.05, Cohen’s *d* = 7.06, healthy level in dark grey). The relationship between SST interneuron inhibition reduction and baseline Pyr neuron firing rate was approximately linear. **e.** Increased interneuron baseline firing rates in depression microcircuit models (*p* < 0.05).

### Reduced SNR of cortical processing in human depression microcircuits

To better understand the implications of the increased baseline firing in depression microcircuits, we compared their evoked response activity to that of healthy microcircuits. We modelled healthy evoked response to a brief stimulus by reproducing the temporal profile and average firing rates in the neuron types as measured in cortical layer 2/3 *in vivo* (in rodents). VIP and PV interneurons were stimulated earliest and consequently silenced SST interneurons. Pyr neurons were stimulated shortly after and responded with a brief peak firing followed by a sustained lower response rate, although still above baseline, that lasted ~100 ms (**Fig. 4a, b**). Over the 5 - 55 ms window post stimulus, Pyr neurons fired at 2.49 ± 0.61 Hz on average (n = 200 randomized microcircuits). We then applied this stimulus paradigm to the depression microcircuits and found no significant change in average Pyr response rate (**Fig. 4c**, 2.52 ± 0.47 Hz, Cohen’s *d* = 0.06). The evoked response of interneurons was similar between healthy and depression microcircuits as well (**Fig. 4i**, SST: 2.74 ± 1.34 vs. 3.21 ± 1.55 Hz, Cohen’s *d* = 0.32; PV: 30.62 ± 4.79 vs. 32.06 ± 3.43 Hz, *d* = 0.35; VIP: 32.88 ± 2.92 vs. 34.94 ± 2.52 Hz, *d* = 0.76).

**Figure 4.**
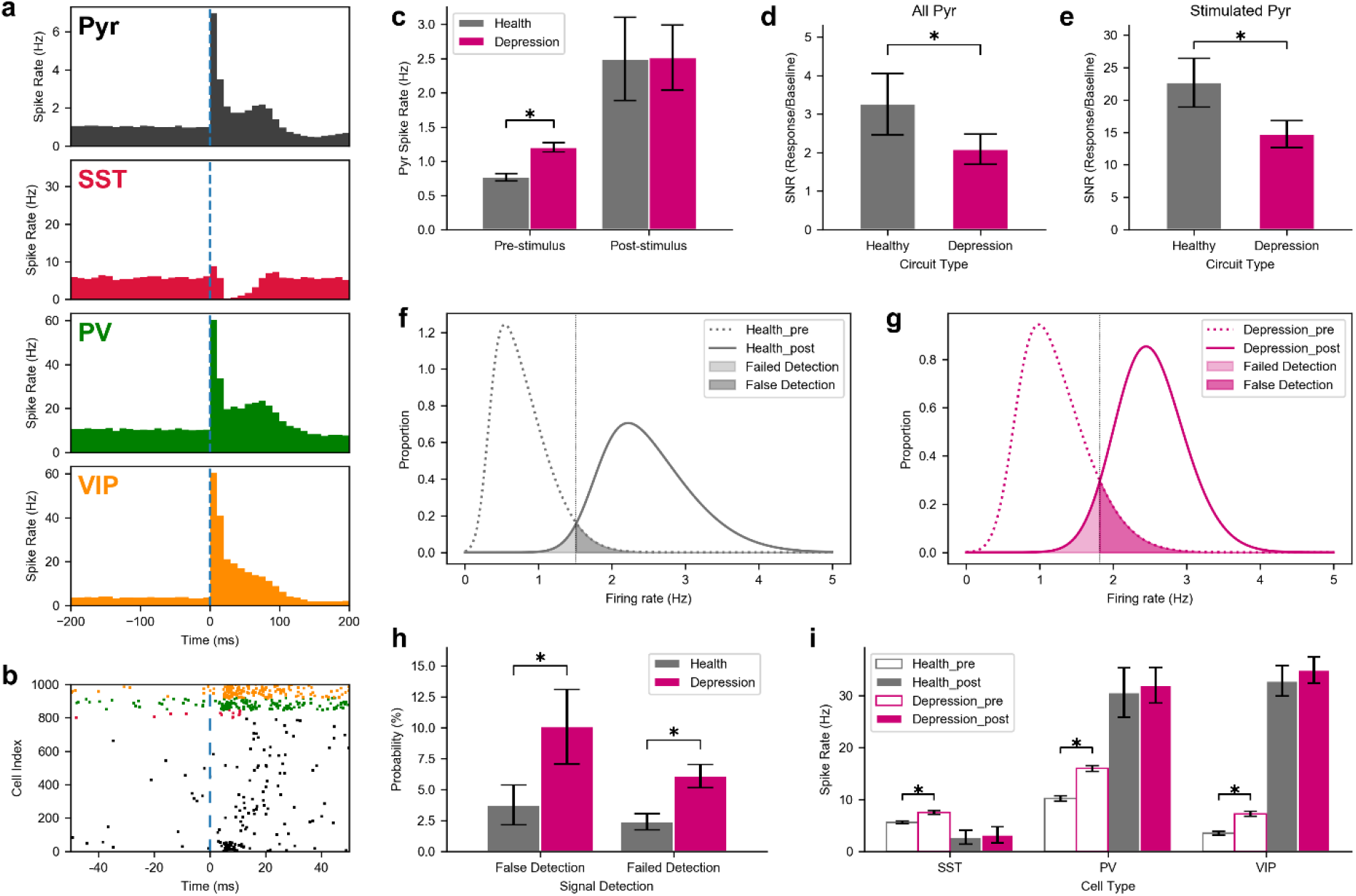
Decreased cortical SNR and impaired stimulus detection in depression microcircuit models. **a.** Average peristimulus time histogram (n = 200 randomized microcircuits) of simulated response to thalamic stimulus in healthy microcircuits reproduce response firing rates and profiles recorded in the different neuron types in awake rodents. Dashed line denotes the stimulus time. **b.** Example raster plot of simulated spike response in a healthy microcircuit model. **c.** Baseline Pyr neuron firing rates in depression microcircuits were significantly higher than in healthy microcircuits (*p* < 0.05, Cohen’s *d* = 7.06), but response rates were similar (Cohen’s *d* = 0.06). **d.** Decreased SNR (ratio of response vs. baseline Pyr neuron firing rates) in depression microcircuits. **e.** Same as d, but for the stimulated Pyr neurons only. **f.** Distribution of pre- and post-stimulus firing rates in 50 ms windows in healthy microcircuits (n = 1950 windows x 200 microcircuits pre-stimulus, n = 200 windows post-stimulus). The vertical dotted line denotes the decision threshold of signal detection (see Methods). **g.** Same as f, but for depression microcircuits. **h.** Increased probability of false detection and failed detection in depression vs. healthy microcircuits. **i.** Baseline and response firing rates in interneurons in healthy and depression microcircuits. Asterisk in all plots denotes *p* < 0.05.

We calculated the overall SNR in the healthy and depression microcircuits, as the ratio of the response activity (signal) and baseline activity (noise) across the Pyr neuronal population, and found significantly (> 50%) reduced SNR in depression microcircuits (**Fig. 4d**, health: 3.26 ± 0.80, depression: 2.09 ± 0.39, *p* < 0.05, Cohen’s *d* = 1.86). The reduction in SNR was similarly significant when examining only the stimulated Pyr neurons (**Fig. 4e**, health: 22.67 ± 3.78, depression: 14.75 ± 2.10, *p* < 0.05, Cohen’s *d* = 2.59). Reducing SST interneuron synaptic or tonic inhibition separately had a similar effect on SNR (synaptic only: 2.56 ± 0.57; tonic only: 2.65 ± 0.57). The effect of the joint synaptic and tonic inhibition reductions was therefore approximately the sum of the separate effects (separate: 1.95 ± 0.80, joint: 2.09 ± 0.39, Cohen’s *d* = 0.22). To further investigate the effect of reduced SNR in cortical processing, we determined the corresponding change in false/failed signal detection rates. We calculated the distribution of spike rates of all Pyr neurons in 50 ms windows pre and post stimulus to calculate the probability of false positive/negative errors in stimulus processing (**Fig. 4f, g**). We found a significant increase, more than doubling, in the false detection rates (health: 3.76 ± 1.60 %, depression: 10.07 ± 3.02 %, *p* < 0.05, Cohen’s *d* = 2.61) and the failed detection rates (health: 2.40 ± 0.67 %, depression: 6.01 ± 0.94 %, *p* < 0.05, Cohen’s *d* = 4.42) in depression (**Fig. 4h**). Reducing SST interneuron synaptic or tonic inhibition separately had a similar effect on the false detection rate (synaptic only: 6.46 ± 2.23 %; tonic only: 5.58 ± 1.94 %) and failed detection rate (synaptic only: 4.10 ± 0.88 %; tonic only: 4.34 ± 0.85%). The effect of the joint synaptic and tonic reductions was therefore approximately the sum of the separate effects on false detection (separate: 8.28 ± 2.96 %, joint: 10.07 ± 3.02 %, Cohen’s *d* = 0.60) and failed detection (separate: 6.04 ± 1.22 %, joint: 6.01 ± 0.94 %, Cohen’s *d* = 0.03).

To determine the effect of SST interneuron inhibition reduction on the processing of top-down inputs, we repeated our previous analysis but with apical dendritic stimulation (corresponding to cortico-cortical inputs). Similarly to basal dendritic stimulation, there was no significant change in response rates (**Fig. 5b**, healthy: 2.96 ± 0.71 Hz, depression: 2.86 ± 0.56 Hz, Cohen’s *d* = 0.16), and there was a similar level of decrease in SNR (**Fig. 5c**, health: 3.88 ± 0.95, depression: 2.39 ± 0.49, *p* < 0.05, Cohen’s *d* = 1.97), and increase in false detection rate (**Fig. 5d**, health: 2.09 ± 0.79 %, depression: 6.21 ± 2.20 %, *p* < 0.05, Cohen’s *d* = 2.49) and failed detection rate (**Fig. 5d**, health: 2.54 ± 0.62 %, depression: 4.73 ± 0.90 %, *p* < 0.05, Cohen’s *d* = 2.83).

**Figure 5.**
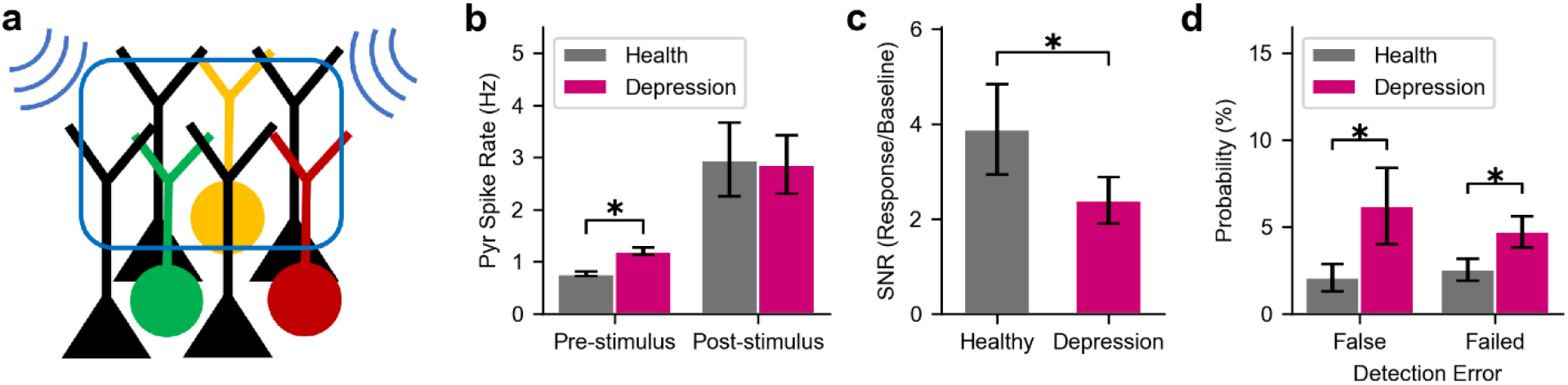
Decreased SNR and increased detection errors in processing top-down inputs. **a.** Schematic of apical dendrite stimulation. **b.** Baseline Pyr neuron firing rates in depression microcircuits were significantly higher than in healthy microcircuits, but response rates were similar. **c.** Decreased SNR (ratio of response vs. baseline Pyr neuron firing rates) in depression microcircuits. **d.** Increased probability of false detection and failed detection in depression vs. healthy microcircuits. Asterisk in all plots denotes *p* < 0.05.

To systematically assess the impact of different levels of SST interneuron inhibition reduction on stimulus processing, we simulated microcircuits with 0 - 100% reduction compared to the healthy level. SNR decreased with the level of SST interneuron inhibition reduction (**Fig. 6a**, 0%: 3.26 ± 0.80, 40%: 2.09 ± 0.39, 100%: 1.30 ± 0.27), with a sharper decrease up to the reduction level estimated in depression (40%), and a more moderate change beyond that level. Interestingly, the rate of false signal detection increased nonlinearly with the level of inhibition reduction (**Fig. 6b**). There was an effect threshold around the level of reduction estimated in depression, with a negligible change for a smaller reduction (0%: 3.76 ± 1.60%; 20%: 4.94 ± 1.95 %, Cohen’s *d* = 0.66) and a sharper increase at the depression level and beyond (40%: 10.07 ± 3.02%, *p* < 0.05, Cohen’s *d* = 2.61; 100%: 40.07 ± 6.70 %, *p* < 0.05, Cohen’s *d* = 7.45). Similar nonlinear increase and threshold effect were observed for failed detection rates (**Fig. 6c**; 0%: 2.40 ± 0.67 %; 20%: 2.21 ± 0.61 %, Cohen’s *d* = −0.30; 40%: 6.09 ± 0.94 %, *p* < 0.05, Cohen’s *d* = 4.5; 100%: 15.44 ± 3.27 %, *p* < 0.05, Cohen’s *d* = 5.5).

**Figure 6.**
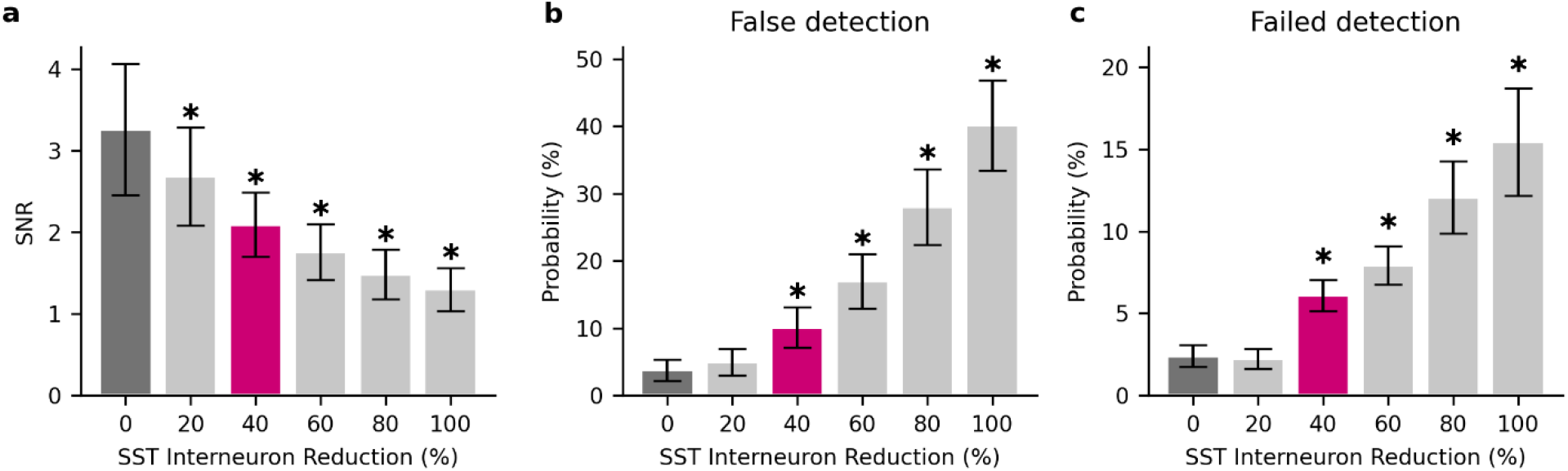
Effect of different levels of reduced SST interneuron inhibition on SNR and signal detection. **a-c.**SNR(a), false signal detection rates(b) and failed signal detection rates(c) at different levels of SST interneuron inhibition reduction. Asterisk in all plots denotes *p* < 0.05 and a large effect size.

## Discussion

This study determined the implications of reduced inhibition in depression on cortical processing in microcircuits, using novel detailed models of human cortical L2/3 microcircuits that integrated human cellular, circuit, and gene-expression data in health and depression. Our simulations of microcircuit response and baseline activity showed that the reduced SST interneuron inhibition in depression estimated by gene expression resulted in a significantly decreased SNR of cortical processing and increased failed/false signal detection rates. Our results thus examined and provided a mechanistic validation for a leading hypothesis of reduced SST interneuron inhibition as a pathological mechanism for dysfunctional processing in depression. This lends support for establishing SST interneuron inhibition as a target mechanism for new pharmacology in treating depression. Furthermore, our results make testable quantitative predictions about the correspondence between different degrees of reduced SST interneuron inhibition and the resulting cognitive deficits that are relevant to symptoms such as rumination and impaired signal detection, which may serve to improve subtyping of depression for better diagnosis, treatment, and drug monitoring in the future.

The reduced SNR and signal detection quality that we found in the depression microcircuit models agrees with previous studies that showed deficits in depression in a variety of tasks involving sensory processing, resulting in impaired detection and decision making^35–37^. Our results may also underlie the inability to break internal loops of thought in rumination symptoms^3^. The increase of failed detection in our depression microcircuit models was of a similar level to a previous study involving a visual attention task^36^, and this study similarly found a larger increase in false detection rate compared to failed detection rate in depression. Other studies that used the Paced Visual Serial Addition Test and Inspection Time Test found that depression patients had slower visual processing speeds and made more calculation errors compared to healthy controls^37,38^. Depression patients have also been shown to perform worse on visual processing tasks such as the Stroop Colour Word Test and in visual surround motion suppression^39,40^. Our demonstration of similar effects on cortical processing of apical (cortico-cortical inputs) and the canonical nature of the cortical microcircuitry^41^ also generalize the implications of our results to higher-order deficits^42^, an important topic that can be explored carefully in future studies. The relationship between dysfunctional microcircuit processing and reduction of SST interneuron inhibition that we quantified may serve as biomarkers for diagnosing depression type and severity^43^. Future studies should examine how the relationship between reduced SST interneuron inhibition and processing is further affected by a disruption in neuromodulation by serotonin and acetylcholine, which is implicated in depression^44–46^. The link between altered inhibition and brain deficits is further supported by studies of other psychiatric disorders^47^. A previous study on the effect of disinhibition in schizophrenia showed a non-linear effect with false signal detection^48^ similar to what we have found. While additional inhibitory mechanisms may be involved, e.g. PV interneurons dysfunction in schizophrenia^49^, this similarity may account for common symptoms relating to mood and processing deficits across different disorders^50^.

The increased baseline firing rate across cell types in the depression microcircuits ultimately resulted from the net effect of different circuit interactions after reducing SST interneuron inhibition, but a few contributing factors can be delineated. The increased Pyr neuron firing led to increase in interneuron firing rates, and the differences in rate change in different interneuron types would partly arise from the differences in synaptic and tonic inhibition reduction from SST interneurons. In addition, the lower level of rate change in SST interneurons would result from being further inhibited by increased inhibition from VIP and PV interneurons. The new excitation/inhibition balance in the depression microcircuits would be restored by the increased inhibition from PV interneurons^51^, but also by the remaining inhibition from SST interneurons.

Lastly, increased activity in VIP interneurons would keep the microcircuit from being overinhibited by the PV and SST interneurons.

We have applied gene expression data to estimate a reduction in the strength (conductance) of inhibitory connections from SST interneurons^11^. SST receptors are colocalized with GABA receptors, and SST is co-released with GABA and strengthens GABA effects through pre and post synaptic mechanisms^52^. Studies have shown that mice lacking SST peptides have altered GABAergic genes in SST neurons that are consistent with reduced GABA function of these cells^53^. Administered SST has also been shown to have antidepressant effects, and reducing GABA in SST interneurons has been shown to produce anxiety behaviour^54,55^. We thus applied the reduction in SST as a reduced GABAergic inhibitory conductance. While the abnormal gene expression may alternatively reflect a reduced number of synapses^56^, the net inhibition decrease in either case would be largely similar. A comparison between conductance decrease and synaptic loss will be interesting to investigate in future studies. We modelled tonic inhibition as a steady-state inhibitory current that is independent of interneuron firing rates, using currently available models. While tonic inhibition depends on interneuron firing rate in addition to the GABA release per synaptic event, the rate dependence remains to be characterized experimentally and modeled, and the timescale is often long enough for tonic inhibition to be considered more steady-state inhibition^57^. Our simulations therefore tested the effects of reduced tonic inhibition directly due to reduced synaptic inhibition and GABA release per synaptic event, together with the rate-dependent effects of reduced synaptic inhibition itself. Rate-dependence of tonic inhibition effects can refine our results when data and models become available.

The effect of synaptic vs. tonic inhibition reduction on cortical processing was similar, and these results are informative for pharmacology treatments that target the SST interneuron inhibition. Recent studies enhanced α5-GABA_A_ receptor function through positive-allosteric modulators to improve SST interneuron signalling, which led to pro-cognitive and anxiolytic effects^16,58^. However, the particular effects of these modulators on cortical circuits are still uncharacterized due to the complex interactions between cell types and differential expression of α5-GABA_A_ receptors in Pyr, SST, and PV neurons^59,60^. Our simulations suggest that reduced synaptic vs. tonic inhibition contribute similarly to the deficits, thus implying both types of inhibition need to be rescued. Therefore, α5-GABA_A_ receptors, which are localized both synaptically and extrasynaptically, serve as good treatment targets^60,61^. Inhibiting VIP interneurons may be another treatment target, as their disproportional increase in firing rate would further exacerbate the reduced contribution of SST interneurons. Our models can serve to test novel treatments systematically *in silico*, and also further optimize and discover new treatment targets.

While our study focused on a systematic characterization of inhibition effects, future studies may also expand the analysis to include excitatory dysfunction in depression, e.g. due to synaptic loss^62^. Our models of depression examined the implication of reduced inhibition at the microcircuit scale, but future studies could explore the implications on multiregional processing in depression, by modelling and stimulating several detailed microcircuits corresponding to the different relevant brain areas and their interactions^63^. Multiregional simulations will help distinguish the similarities and differences of SST interneuron inhibition effects in various neurological disorders where these interneurons are implicated^9,16^. Multiregional processing simulations will also enable examining the implications on the later stages of the cortical response, where SST interneurons may provide feedback inhibition to reduce excitation when stimuli are similar to contextual surroundings^64^, or serve to shorten response duration^65^. Pathologically reduced inhibition in such cases may result in prolonged response time and thus slower processing of subsequent stimuli^43^.

We developed novel models of human cortical microcircuits to study depression in the context of the human cortex. Our microcircuit models were constrained with circuit data from different regions due to availability, but aimed primarily at representing a canonical cortical microcircuit and response to a brief stimulus. While different regions such as sensory and association regions have some differences in wiring and function when considering all six layers of cortex, the canonical L2/3 microcircuitry and the response to brief stimulus are mostly similar across regions^41^. We were able to improve upon existing models of human L2/3 neurons to better reproduce the firing frequency-input relationship and the sag voltage, thus capturing important input integration properties of neurons within the microcircuit. Further improvements to our models can include dendritic ion-channels mediating backpropagating action potentials and calcium spikes that have been well characterized in rodents^29^. These dendritic properties still remain to be better characterized in humans, but recent studies suggest that calcium spikes may play a role in mediating nonlinear processing in human Pyr neurons^22^. We note, however, that our microcircuit models are still capable of NMDA nonlinearities, which play an important role in the cortical microcircuit^66^. A further refinement of the models would be constraining the connectivity between neuron types in the human cortex, which is largely unavailable at present but is expected to become available in the near future^67^. Nevertheless, the model spike oscillations in theta and alpha frequency bands, and the 1/f relationship between power and log frequency agree with the oscillations seen in human cortical signals experimentally^31–34^ and serve as a validation in support of the models capturing key properties of human cortical L2/3 microcircuits. As human data from other cortical layers becomes more available, future directions should examine the implication of reduced SST interneuron inhibition in depression on signal propagation across layers^68^. We examined the effect of SST interneuron inhibition on cortical processing using the response rate profile in the different neuron types based on available somatosensory studies in rodents, due to the lack of similar data in humans. The relevance of rodent response data is supported by the important similarities between rodent and human cortex in terms of ion-channel mechanisms, neuron types, and microcircuit connection motifs such as the SST interneuron inhibitory disynaptic loop^14,15,69^. Future studies should investigate how the effects we found differ in rodent microcircuit models, with smaller neurons and weaker synapses, to better establish the preclinical relevance of the rodent animal model of depression. Due to the above-mentioned species similarities in neuron types and circuit motifs we expect the results to largely hold for rodent microcircuits as well, although with possible nuanced differences in the extent and magnitude of the effects. Our detailed models of healthy microcircuits can further serve to study the cellular and circuit mechanisms of other neurological disorders such as schizophrenia and epilepsy. Furthermore, our models are implemented in LFPy^70^ and thus can be used to study the associated electroencephalography signals and local field potentials from our depression microcircuit models to identify biomarkers for improving diagnosis and monitoring^71^.

## Methods

### Experimental Data

We used whole-cell recordings from human medial temporal gyrus, L2/3 Pyr neurons^24^ (n = 28 neurons, 9 neurons from 3 male subjects and 19 neurons from 5 female subjects), and putative SST (Neuron ID: 571700636), PV (Neuron ID: 529807751) and VIP (Neuron ID: 525018757) interneurons available from the Allen Brain Atlas^72,73^. We used reconstructed human neuron morphologies from the Allen Brain Atlas for a Pyr neuron (Neuron ID: 531526539) and the above three interneurons. We used five hyperpolarizing and depolarizing current steps from the electrophysiological data for model optimizations (**Tables S2 - S5**). Three depolarizing supra-threshold current steps were used for targeting appropriate firing features at low, medium and high firing rates. A small hyperpolarizing step current was used for fitting passive features, and a large hyperpolarizing current step was used for fitting sag voltage.

In addition, we used current recordings of tonic inhibition in human cortical L2/3 Pyr neurons (10 cells: 9 cells from 3 male subjects, 1 cell from 1 female subject). Written informed consent was obtained from all participants, in accordance with the Declaration of Helsinki and the University Health Network Research Ethics board. The data was collected using surgery resection, solutions, tissue preparation, and recording equipment described previously^24^. For voltage-clamp recordings of tonic current, low-resistance patch pipettes (2 – 4 MΩ) were filled with a CsCl-based solution containing (in mM) 140 CsCl, 10 EGTA, 10 Hepes, 2 MgCl2, 2 Na2ATP, 0.3 GTP, and 5 QX314 adjusted to pH 7.3 with CsOH. Experiments were performed with excitatory (APV 25 μM, Sigma; CNQX 10 μM, Sigma) and GABA_B_ (CGP-35348 10 μM, Sigma) synaptic activity blocked as in previous studies^74,75^. The junction potential was calculated to be 4.3 mV and the holding potential was −74.3 mV after junction potential correction. Mean amplitude of baseline tonic current in presence of GABA (5 μM) and glutamatergic blocker was 65.8 ± 10.2 pA (n = 10 cells).

### Human L2/3 Microcircuit Models

We simulated cortical L2/3 microcircuits comprised of 1000 neurons distributed along a 500×500×950 μm^3^ volume (250 to 1200 μm below pia, spanning L2/3^21^) using NEURON^76^ and LFPy^70^. We used human data to constrain the models where available (**Table S1** and see below). Neurons of a given type had the same model (morphology and biophysical properties, see below) but differed in synaptic connectivity and background input (see below). The proportions of the four neuron types in the microcircuit were: 80% Pyr, 5% SST, 7% PV, and 8% VIP, according to the relative neuron densities found in the rodent and human cortical L2/3 in previous studies^77,78^, and RNA-seq data from the Allen Human Brain Atlas^17,73^.

### Neuron Models

We derived multi-compartmental conductance-based models for the different neuron types using multi-objective optimization with a genetic algorithm from previous in-house code^29^ for optimizing the Pyr and SST neuron models, and the BluePyOpt Python module^79^ for optimizing the PV and VIP neuron models. We used a set of ion channel mechanisms taken unchanged from previously published models^29,80^. Pyr and SST neuron models were fitted using a two-step process, where first h-current and passive parameters (*ḡ_H_*, *ḡ_Leak_,*) were fitted to capture passive and sag voltage features during hyperpolarizing current steps, and then other ion channel parameters (*ḡ_Na_*, *ḡ_K_*, *ḡ_Ca_*) were fitted to capture spiking features during depolarizing current steps (**Tables S2 - S3**). The PV and VIP neuron models were fitted in one step where passive and firing features were optimized simultaneously (**Tables S4 - S5**). For all models, *R_a_* = 100 Ω cm, *E_Na_* = 50 mV, *E_K_*= −85 mV, and *CaDynamics_gamma_* = 0.0005^29^. Axonal NaT kinetics parameters were *Vshift_m_* = 0, *Vshift_h_* = 10, *Slope_m_* = 9, and *Slope_h_* = 6 for all models^29,80^. For somatic Na_T_ kinetics, *Vshift_m_* = 13, *Vshift_h_* = 15, *Slope_m_* = 7, and *Slope_h_* = 6^29,80^, except for the PV neuron model where these parameters at the soma were the same as in the axon to compensate for reduced action potential propagation brought on by the axon replacement method. The specific membrane capacitance (*c_m_*) was 1 μF/cm^2^, and 2 μF/cm^2^ in the dendrites to compensate for spine area loss in Pyr neuron reconstructions^81,82^ or where required to reproduce membrane time constants in interneurons, possibly due to errors in dendritic diameter estimation (**Table S6**). *ḡ_H_* was distributed uniformly across all dendritic sections, with the exception of Pyr apical dendrites which followed an exponential increase with distance from soma as follows:

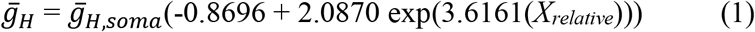

where *X_relative_* is the relative distance from soma to maximal length (i.e. from 0 to 1)^29^. Model optimization was run using parallel computing clusters (SciNet^83,84^: 400 processors with a population size of 1000, for 1000 generations and an approximate total runtime of 2 hours; Neuroscience Gateway, NSG^85^: 400 processors with a population size of 400, for 300 generations and an approximate total runtime of 5 hours). Model performance in reproducing the electrophysiological features was assessed in terms of standard deviation from the experimental mean. For Pyr neuron models we used the statistics over the set of recorded neurons (see above). For the interneurons, where only a single human neuron was clearly identified for each neuron type, we used the values of the human neuron with population variance from the rodent literature^26–28^.

### Synaptic Connectivity Models

Where possible, we constrained synaptic parameters using the human experimental literature. Otherwise, we used parameters from the rodent literature (the Blue Brain Project^86^). We used previous models of NMDA/AMPA excitatory and GABA_A_ inhibitory synapses, that incorporated presynaptic short-term plasticity parameters for vesicle-usage, facilitation, and depression, as well as separate time constant parameters for the AMPA and NMDA components of excitatory synapses^66,71,87^. We set the same time constant parameters for all connection types (*τ_rise,NMDA_* = 2 ms; *τ_decay,NMDA_* = 65 ms; *τ_rise,AMPA_* = 0.3 ms; *τ_decay,AMPA_* = 3 ms; *τ_rise,GABA_* = 1 ms; *τ_decay,GABA_* = 10 ms), as well as the reversal potential values (*E_exc_* = 0 mV; *E_inh_* = −80 mV).

We fitted the synaptic conductance, and vesicle usage parameters^66^ where available, using the human experimental literature for most of the key connection types: Pyr→Pyr connections^20^, Pyr→SST connections and SST→Pyr connections^14^, PV→Pyr connections^19^, and Pyr→PV connections^88^. We simulated the experimental conditions (e.g., chloride reversal potential, and holding currents) and adjusted the conductance and vesicle usage parameters to achieve the target postsynaptic amplitudes and failure rates on average across 200 randomizations of synaptic locations and events. For the conductance and vesicle usage parameters of all other connection types, as well as the depression, facilitation, and numbers of synaptic contacts for all connections, we used model parameters reported for rodents by the Blue Brain Project^86,89^. The synaptic parameters of the different connections are summarized in **Table S7**. Synaptic locations onto Pyr neurons were random but depended on the connection type, where Pyr→Pyr synapses were placed on both basal and apical dendritic compartments, PV→Pyr connections were placed on basal dendritic compartments, and SST→Pyr connections were placed on apical dendritic compartments. Apart from these specifications, synapse locations were chosen randomly from a uniform distribution.

We set unidirectional connection probability (*p_con_*) according to the rodent literature (Blue Brain Project and Allen Brain Institute), except for Pyr→Pyr connections, where *p_con_* was set to 15% according to the human literature^20^. We adjusted some connection probabilities guided by the reported experimental ranges to reproduce the intrinsic activity in the microcircuit (see below). The connection probabilities in the microcircuit are summarized in **Table S7**.

### Tonic Inhibition Models

We modelled tonic inhibition using a model for outward rectifying tonic inhibition^90^ and estimated the conductance of tonic inhibition (*G_tonic_*; uniformly across all somatic, basal, and apical compartments) using current magnitude in human cortical L2/3 Pyr neurons (see above), which agreed with values from the human layer 5/6 literature^30^. We replicated the experimental conditions by setting the GABA reversal potential to −5 mV (i.e. consistent with a high chloride intracellular solution), setting the holding potential to −75 mV in voltage-clamp mode, and reproduced the target tonic inhibition current amplitude with G_tonic_: 0.938 mS/cm^2^. We used the same value for the interneurons since the total tonic inhibition current recorded in interneurons was similar to that of Pyr neurons after correcting for cell capacitance^30^.

### Modelling Microcircuit Baseline and Response Activity

We constrained the microcircuit to generate spike rates at baseline and during response (see below) as reported for the different neuron types *in vivo*. For baseline activity of Pyr neurons, we used values recorded in humans and monkeys *in vivo*^91^ (Subject 1: 0.66 ± 0.51 Hz; Subject 2: 0.32 ± 0.38 Hz; Monkey: 1.31 ± 1.11 Hz), and for baseline activity of the interneuron types we used values recorded in rodents *in vivo* (SST: 6.3 ± 0.6 Hz; PV: 9.4 ± 2.1 Hz; VIP: 3.7 ± 0.7 Hz)^1,65^. We note that baseline Pyr rates in human and rodents in the above studies were largely similar. We reproduced the baseline firing rates by adjusting the connection probability values guided by the reported experimental ranges (see above) for all connection types except Pyr→Pyr connections, and by adjusting the background input (see below). We calculated the simulated rates across non-silent neurons (> 0.2 Hz), as in the above experimental studies. We further constrained the microcircuit model to reproduce a doubling of baseline Pyr neuron spike rates when SST interneurons were silenced, as seen experimentally^1^ (baseline: 1.2 ± 0.2 Hz; SST interneurons silenced: 2.2 ± 0.3 Hz).

The microcircuit received random background excitatory input using Ornstein-Uhlenbeck (OU) point processes^92^, placed at halfway the length of each dendritic arbor to ensure similar levels of inputs along each dendritic path to the soma. For the Pyr neuron models, we placed 5 additional OU processes along the apical trunk at 10%, 30%, 50%, 70%, and 90% of the apical dendritic length. We set the base excitatory OU conductances to the following: Pyr = 28 pS; SST = 30 pS; PV = 280 pS; VIP = 66 pS. We set the inhibitory OU conductance to 0, since the model microcircuit provided sufficient inhibition. Furthermore, we scaled the OU conductance values to increase with distance from soma by multiplying them with the exponent of the relative distance from soma (ranging from 0 to 1): *ḡ_OU_ = ḡ* × *exp*(*X_relative_*).

We modelled the spike response rates and profiles using values recorded in the different neuron types *in vivo*, in behaving rodents^1^ (change in firing rate compared to pre-stimulus rate: Pyr 2.2 ± 1.0 Hz, SST: −2.9 ± 0.9 Hz; PV: 21.8 ± 8.9 Hz; VIP: 14.0 ± 3.0 Hz). We reproduced the response rates by synaptic stimulation of the model microcircuits representing a bottom-up (thalamic) input. We tuned the microcircuit connectivity, the stimulus input conductance and stimulus timing in the different neuron types to reproduce the experimental response profiles and timing of activation^1,65^. The model neurons were stimulated using excitatory AMPA/NMDA synapses with the same synaptic dynamics and number of contacts as the cortical excitatory synapses above. 55 Pyr neurons were stimulated in the basal dendrites, with 2 – 4 ms delay post-stimulus and a conductance of 4 nS. 35 PV interneurons were stimulated with a delay of 2 - 2.5 ms and a conductance of 2 nS. VIP interneurons were stimulated in two groups and phases: early (65 VIP interneurons, delay = 0.5 - 4.5 ms, conductance = 2.8 nS) and late (80 VIP interneurons, delay = 7 - 12 ms, conductance = 2.2 nS). This stimulus paradigm was applied to 200 randomized healthy and depression microcircuits. Average pre-stimulus rates were calculated over the 2 seconds before stimulus onset. Average post-stimulus rates were calculated over the 5 - 55 ms window after stimulus onset. To examine and visualize the stimulus response in each of the four cell types, we calculated peristimulus time histograms of the spike counts of non-silent cells (> 0.2 Hz) in 10 ms bins, 200 ms before and after stimulus, pooled across 200 randomized microcircuits. We also tested apical dendritic stimulation, with parameters as above except that the input synapses were distributed on the apical dendrites instead of the basal dendrites, and the stimulated Pyr neuron population size was 85 neurons.

### Spikes PSD

Power spectral density (PSD) of Pyr neuron population spiking during baseline microcircuit activity was computed by first converting the spike times into binary spike train vectors and then summing the binary spike train vectors across all Pyr neurons. PSD was then computed from the summed spike train vectors using Welch’s method^93,94^ from the SciPy python module (nperseg=100,000 sampling points, equivalent to 2.5s time windows). PSD vectors across random seeds were then bootstrapped at each frequency 500 times, from which the resulting bootstrapped means and 95% confidence intervals were computed.

### Human Depression Microcircuit Models

We modelled depression microcircuits by reducing the conductance of SST interneuron synaptic and tonic inhibition on all cell types by 40%, as indicated by the 40% reduction of SST expression in SST interneurons in L2/3 in post-mortem brain tissue of patients with depression^11^. For Pyr neurons, we selectively decreased the tonic inhibition by 40% in the apical dendrites. For each interneuron, we estimated the relative contribution of SST interneurons to the total inhibitory input and reduced the tonic inhibition by 40% of the relative contribution. This relative contribution was calculated by multiplying the SST synaptic conductance, connection probability, and the number of contacts, and dividing by the summed contributions of all interneuron types.

### False/Failed Signal Detection Rates

We calculated the error rates in stimulus processing by first computing the distribution of pre-stimulus firing rates of all Pyr neurons per circuit run (sliding windows of 50 ms over 2 seconds pre-stimulus, windows sliding in 1 ms steps for a total of 1950 windows), and the distribution of post-stimulus spike rates of all Pyr neurons (in the 5 – 55 ms window post stimulus) across 200 randomized microcircuits. The intersection point between the two distributions was set as the stimulus detection threshold. The probability of false detection was calculated by the integral of the pre-stimulus distribution above the detection threshold divided by the integral of the entire pre-stimulus distribution. The probability of failed detection was calculated by the integral of the post-stimulus distribution below the detection threshold divided by the integral of the entire post-stimulus distribution.

### Statistical Tests

We determined statistical significance using paired-sample or two-sample t-test, where appropriate. We calculated the effect size using Cohen’s *d* (the difference in means divided by the pooled standard deviation).

### Model and code availability

All models and simulation code will be available openly online (ModelDB, accession number: TBA).

## Acknowledgements

HKY, AGM, FM and EH thank the Krembil Foundation for funding support. HKY and EH were also supported by a stipend award from the Department of Physiology at University of Toronto.

